# The Genome of BAM-degrading *Aminobacter* sp. MSH1 with Several Low Copy Plasmids

**DOI:** 10.1101/307967

**Authors:** Tue Kjærgaard Nielsen, Ole Hylling, Lea Ellegaard-Jensen, Jens Aamand, Lars Hestbjerg Hansen

## Abstract

As one of the only described degraders of the recalcitrant metabolite 2,6-dichlorobenzamide (BAM) of the pesticide dichlobenil, *Aminobacter* sp. MSH1 has been intensively studied for its characteristics with regards to physiology and its use in bioremediation. Two plasmid sequences from strain MSH1 have previously been published, while the remaining genome sequence has been left uninvestigated. We here present the complete genome sequence of this important strain, which consists of a chromosome, two megaplasmids and five smaller plasmids. Intriguingly, the plasmid copy numbers are mostly below one per bacterial chromosome, indicating that plasmids in strain MSH1 are under very unstable conservation. The results of this report improve our understanding of the genomic dynamics of *Aminobacter* sp. MSH1.

## Introduction

*Aminobacter sp*. MSH1 is a Gram-negative, motile rod isolated from a plant nursery courtyard soil, previously exposed to the pesticide dichlobenil (Sørensen et al. 2007). *Aminobacter* MSH1 and two other *Aminobacter* strains (ASI1 and ASI2) are the only bacteria reported in the literature capable of utilizing the recalcitrant dichlobenil metabolite 2,6-dichlorobenzamide (BAM) as their sole carbon source and mineralize it at nanomolar concentrations (Simonsen et al. 2006; Sørensen et al. 2007). This characteristic is of immense interest since BAM is among the most common micro-pollutant found in groundwater aquifers of several European countries (Porazzi et al. 2005; Törnquist et al. 2007; Schipper et al. 2008; Pukkila and Kontro 2014; Vandermaesen et al. 2016), often exceeding drinking water threshold limits set by the European Union (0.1 μg/L) (The Council of the European Union 1998).

Since many countries rely partially or solely on groundwater for drinking water production, remediation of pesticide polluted groundwater by bioaugmentation is proposed as a cost-effective and sustainable biotechnological method (Benner et al. 2013). Strain MSH1 has been suggested as a promising candidate to remediate BAM-polluted groundwater, and thus prevent the costly closures of abstraction wells in drinking water production (Björklund et al. 2011). Laboratory and pilot scale studies have applied MSH1 to drinking water treatment sand filter (SF) units for biological removal of BAM contamination (Albers et al. 2015; Ellegaard-Jensen et al. 2016; Horemans, Raes, Vandermaesen, et al. 2017). However, MSH1 is challenged by the oligotrophic environment of SFs that may inhibit degradation and maintenance of the MSH1 population (Ellegaard-Jensen et al. 2017).

Over the past decade, studies of MSH1 have uncovered and described several characteristics, important in this context; BAM-catabolic genes, enzyme characterization, metabolic pathways, adhesion properties, surface colonization and invasion of natural SF communities (Simonsen et al. 2012; Albers et al. 2014; T’Syen et al. 2015; Sekhar et al. 2016; Horemans, Raes, Brocatus, et al. 2017; Horemans, Vandermaesen, et al. 2017). However, despite comprehensive attention, whole genome sequencing of MSH1 is still absent from literature. We report for the first time the complicated complete genome of *Aminobacter sp*. MSH1, which consists of a chromosome, two megaplasmids, and 5 plasmids in the range of 31-97 kb in size.

## Materials and Methods

*Aminobacter* sp. MSH1 was obtained from the strain collection of the laboratory that originally isolated the bacterium. Strain MSH1 was grown in shaking R2B medium for 72 hours at 22°C. 2 ml culture was used for extraction of high molecular weight (HMW) DNA using the MasterPure^™^ DNA Purification Kit (Epicentre, Madison, WI, USA), using the protocol for cell samples. The purity and concentration of extracted DNA were recorded with a NanoDrop 2000c and a Qubit^®^ 2.0 fluorometer (Thermo Fisher Scientific, Walther, MA, USA), respectively. An Illumina Nextera XT library was prepared for paired-end sequencing on an Illumina NextSeq 500 with a Mid Output v2 kit (300 cycles) (Illumina Inc., San Diego, CA, USA). For Oxford Nanopore sequencing, a library was prepared using the Rapid Sequencing kit (SQK-RAD004). This was loaded on an R9.4 flow cell and sequenced using MinKnow (v1.11.5). Nanopore reads were basecalled with albacore (v2.1.10) without quality filtering of reads. Only reads longer than 5,000 bp were retained and sequencing adapters were trimmed using porechop (v0.2.3). For Illumina sequencing, 2×151 bp paired-end reads were trimmed for contaminating adapter sequence and low quality bases (<Q20) in the ends of the reads were removed using Cutadapt (v1.8.3) (Martin 2011). Paired-end reads that overlapped were merged with AdapterRemoval (v2.1.0) (Martin 2011). A hybrid genome assembly with Nanopore and Illumina reads was performed using Unicycler (v0.4.3) (Wick et al. 2017). An automatic annotation was performed with Prokka (v1.11) (Seemann 2014) which was manually curated for plasmids pBAM1 and pBAM2 using BLAST and the associated Conserved Domain Database (Altschul et al. 1990; Marchler-Bauer et al. 2015). Sequence alignment was produced with MUSCLE (v3.0) (Edgar 2008), as implemented in CLC Genomics Workbench (v11) (QIAGEN, Hilden, Germany).

The complete genome assembly of *Aminobacter* sp. MSH1 was deposited in GenBank under accession numbers CP028966-CP028973.

## Results and Discussion

### Complete genome of *Aminobacter* sp. MSH1

Sequencing of *Aminobacter* sp. MSH1 on Illumina NextSeq yielded 8.5M paired-end reads before quality filtering. 2.9M filtered paired-end reads were not overlapping, while 2.7M reads were merged for a total Illumina data size of 661 Mbp. Nanopore sequencing resulted in 287K reads with a total data size of 5.5 Gbp, an average read length of 19 kb, and a read N50 of 33 kb. After trimming of Nanopore adapters and removal of short reads (<5 kb), 213K Nanopore reads with a total of 5.27 Gbp remained for hybrid genome assembly which resulted in eight circular and closed DNA replicons (Figure 1).

**Figure 1.**
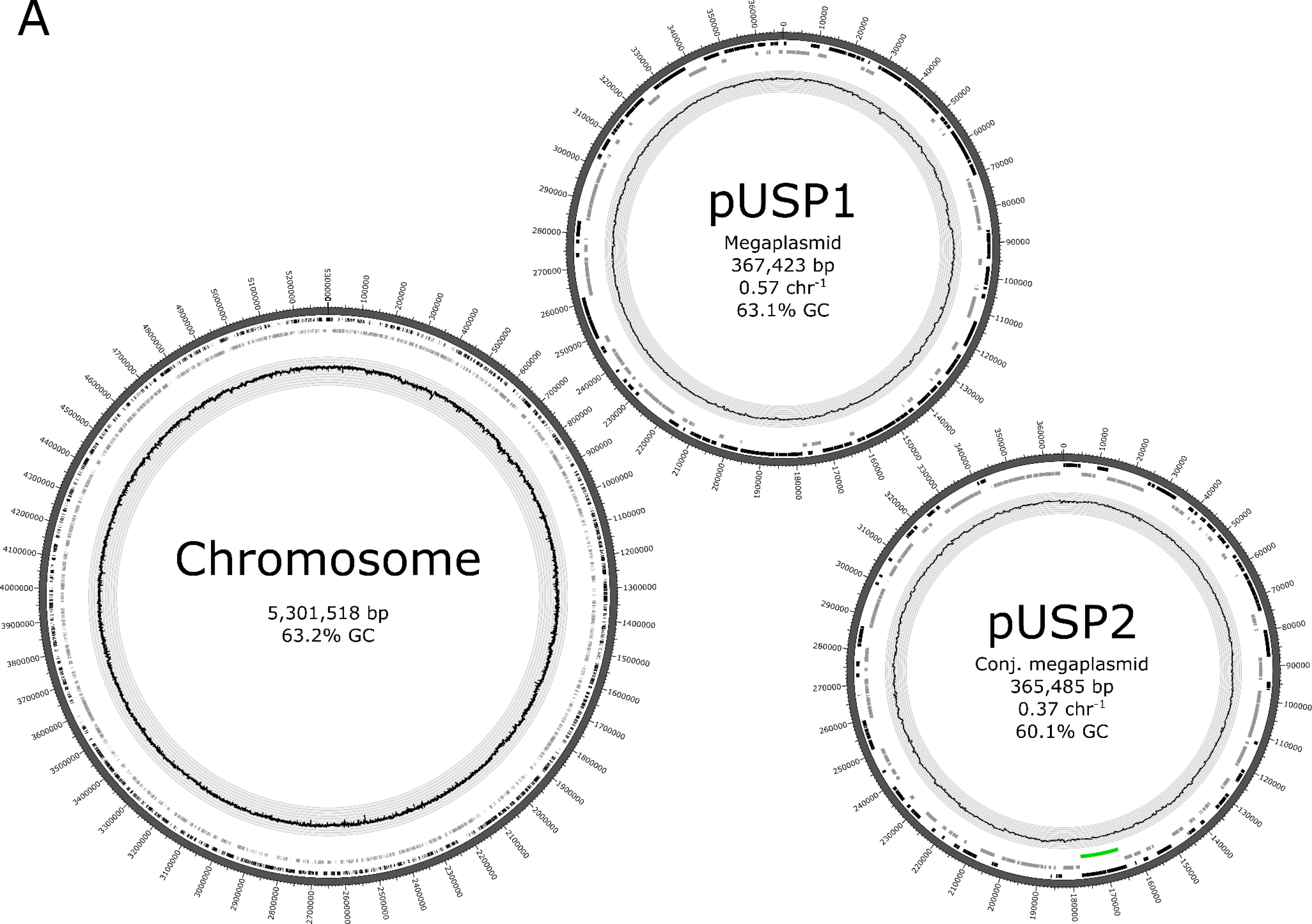
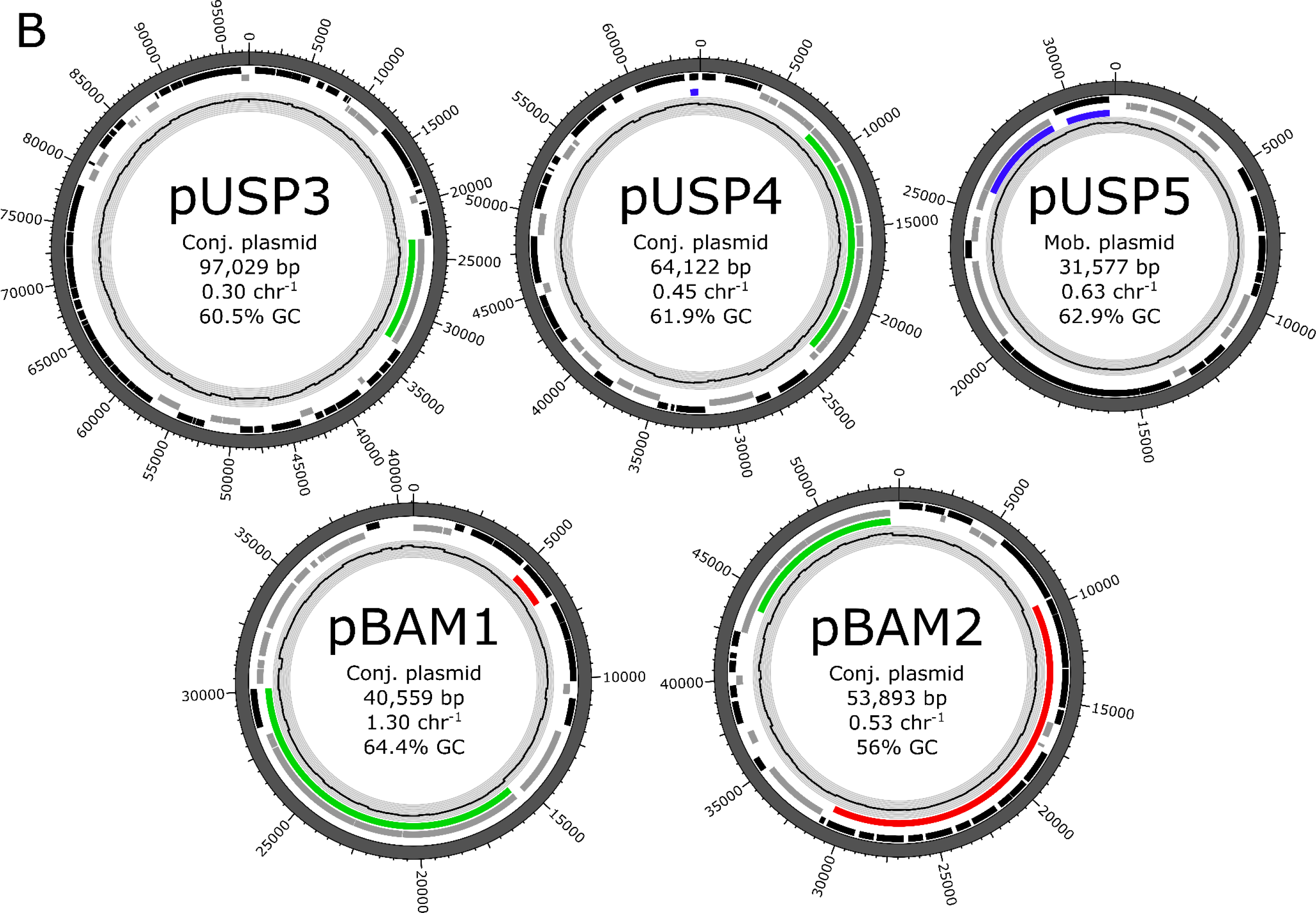
Complete genome assembly of *Aminobacter* sp. MSH1 with eight replicons determined by Unicycler assembler to be complete and circular. The complete genome consists of a chromosome and two megaplasmids (A) and five plasmids ranging in size from 97 kb to 31.6 kb (B). For all replicons, the GC content (window size 500 nt) is shown as a graph with grey background. Genes on forward strands are shown as black bars, while genes on reverse strands are shown as grey bars. Type IV secretion systems for conjugative plasmid transfer are highlighted with green bars, while genes putatively involved in plasmid mobilization are marked with blue bars. The *bbdA* gene on pBAM1 and the *bbd* gene cluster on pBAM2 are highlighted with red bars. The copy number per chromosome of the plasmids are reported by the Unicycler assembler and are based solely on Illumina data.

Megaplasmid pUSP1 has no apparent genes encoding conjugative machinery or plasmid mobilization, suggesting that this replicon might be a possible secondary chromosome. However, the low copy number of 0.57 per chromosome indicates that this replicon is not essential for survival of strain MSH1. The second megaplasmid, pUSP2, does harbor a *vir* operon encoding a type IV secretion system (T4SS) for plasmid conjugation. Furthermore, plasmids pUSP3-4 and pBAM1-2 each have genes encoding putative T4SS proteins, whereas pUSP5 seems to possibly be a mobilizable plasmid due to presence of genes encoding a TraG coupling protein and TraA conjugative transfer relaxase but no T4SS. Plasmids pUSP1-3 and pBAM2 are all *repABC* family plasmids. pBAM1 was previously found to be an IncP1 plasmid through analysis of the plasmid replication initiator gene *trfA* (T’Syen et al. 2015).

Surprisingly, only plasmid pBAM1 had a copy number per chromosome higher than one. It was previously reported that, under non-selective BAM-free conditions, a phenotype of MSH1 incapable of BAM mineralization rapidly increased in abundance. It was speculated to be caused by a loss of plasmid pBAM2, harboring genes responsible for degradation of 2,6-DCBA – the first metabolite of BAM. The stability of BAM mineralization was monitored with qPCR targeting the single copy *bbdA* and *bbdD* genes on pBAM1 and pBAM2, respectively, and calculating their abundance relative to 16S rRNA gene copy numbers. It was found that a culture grown in mineral salt medium with 200 mg/liter BAM as sole carbon source sustained gene copy ratios of *bbdA*/16S and *bbdD*/16S of 2.4 and 2.2, respectively. While the *bbdA*/16S rRNA ratio was sustained under varying incubation settings, the *bbdD*/16S ratio rapidly diminished when other C sources than BAM were available (Horemans, Raes, Brocatus, et al. 2017). The complete genome of strain MSH1 includes two 16S rRNA gene copies on the chromosome. This results in a plasmid copy number of pBAM1 and pBAM2 under BAM selective conditions of 1.2 and 1.1, respectively, based on the previously determined *bbdA*- and *bbdD* to 16S rRNA gene ratios. For pBAM1, this approximately agrees with the observed plasmid copy number based on Illumina data coverage. However, for the remaining six plasmids, the copy number per chromosome is curiously below one, indicating that these plasmids may be unstable in strain MSH1 or may be maintained at unusually low copy numbers compared to the chromosome. Future studies should investigate the unusually low copy numbers per chromosome of the plasmids in MSH1, perhaps owing to a possible chromosomal polyploidy as seen in multiple bacterial lineages (Oliverio and Katz 2014).

### Characteristics of pBAM1

The first step in degradation of BAM is performed by the BbdA amidase which converts it to 2,6-dichlorobenzoic acid (DCBA) (T’Syen et al. 2015). The *bbdA* gene is located on pBAM1 (Figure 1), while two *bbd* gene clusters located on pBAM2 encodes enzymes converting DCBA to intermediate compounds in the Krebs cycle (Horemans, Raes, Brocatus, et al. 2017). Only pBAM1 is maintained in the population at a copy number of more than one per chromosome, suggesting that it is either more stable than the other plasmids or that there are multiple copies of the chromosome per cell.

The gene encoding the pivotal BbdA amidase on pBAM1, is located immediately upstream of two genes encoding a putative IS*5* transposase. While no closely related homologs of BbdA currently exists in the NCBI nr database, the putative IS*5* transposase has 90% nucleotide similarity to sequences from *Comamonadaceae* bacteria, indicating that this transposase has a broad phylogenetic dispersal. The proximity of *bbdA* to a transposase furthermore indicates that this gene has been transferred from an unknown origin to plasmid pBAM1.

### Characteristics of pBAM2

Curiously, a 2,557 bp region on pBAM2, encoding the genes for gluthathione S-transferase (*bbdI*) and gluthathione-disulfide reductase (*bbdJ*), has apparently undergone duplication and occurs in three perfect consecutive repeats followed by one imperfect repeat (Figure 2). These genes are part of the cluster that is responsible for the conversion of 2,6-DCBA to Krebs cycle intermediates, as previously established (Horemans, Raes, Brocatus, et al. 2017). The fourth, imperfect, repeat encompasses the gene previously annotated as the 1,500 bp gene *bbdK* encoding a gluthathione-disulfide reductase. This gene is likely derived from the 1,392 bp *bbdJ* genes occurring in the upstream three perfect repeats, as the first 1,377 bp are identical in *bbdJ* and *bbdK* (results not shown). It is not known whether *bbdK* is a non-functional pseudogene derived from *bbdJ*, but considering the redundancy of genes encoding gluthathione-disulfide reductases on pBAM2, it seems likely that *bbdK* has been subjected to spurious mutations.

**Figure 2.**
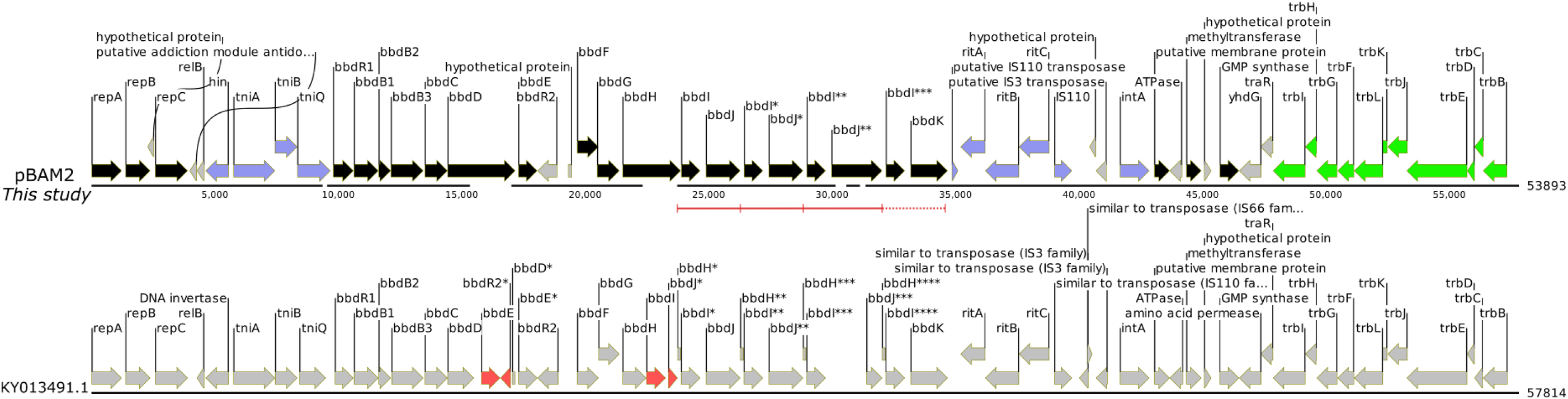
Plasmid pBAM2 from this study aligned with MUSCLE to the already published variant. The alignment is shown as black lines under the genes for both sequences. Notice that gaps are introduced by MUSCLE in the 53,893 bp pBAM2 sequence to align to the previously published 57,814 bp KY013491.1 sequence. For pBAM2 (this study), genes on forward strands are in black, while genes on reverse strands are in grey. Genes encoding putative mobile genetic elements are highlighted in blue, while the T4SS genes are highlighted in green. The four repeats of *bbdIJ* are underscored in red brackets with the fourth, imperfect, repeat shown with a dotted bracket.

The *bbdB-KR* gene region in the presented complete genome assembly shows some discrepancies with the sequence of pBAM2 that was previously deposited in GenBank (accession number KY013491.1). Specifically, the previous pBAM2 assembly is 3,921 bp longer and contains additional copies of the *bbdE, bbdR2, bbdI*, and *bbdJ* genes (Figure 2). The structure of pBAM2, as shown in this study, was verified by mapping of 250 Nanopore reads (minimum length 50,000 bp) to the plasmid sequence, confirming that the pBAM1 assembly is complete. While the MSH1 strain used for sequencing in this study was obtained from the laboratory from which it was originally isolated, it cannot be excluded that genetic rearrangements may have occurred between laboratory strains which have led to the observed discrepancies.

Except for the plasmid backbone (*trb* and *rep* genes), there are no significantly similar sequences in the NCBI nr/nt database (BLASTN (Altschul et al. 1990) search on April 11^th^ 2018), which obstructs any attempts to derive how pBAM2 and the *bbd* gene cluster has evolved. Future studies will hopefully lead to the discovery of sequences related to pBAM2. However, several putative mobile genetic elements are located near the *bbd* gene cluster, including the *ritABC* genes that were recently shown to form circular DNA intermediate molecules as a likely intermediate in transposition (Nielsen et al. 2017). This makes it probable that the *bbd* genes have been transferred from another replicon to pBAM2. Furthermore, pBAM2 has a lower GC content than the remaining genome, indicating that is has been transferred to MSH1 from another bacterium. Whereas BAM mineralization occur only rarely in environmental samples, 2,6-DCBA mineralization is much more common (Vandermaesen et al. 2016), supporting that the *bbdB-KR* genes are more widely spread than the *bbdA* gene.

